# Ebselen as a highly active inhibitor of PL^Pro^CoV2

**DOI:** 10.1101/2020.05.17.100768

**Authors:** Ewelina Węglarz-Tomczak, Jakub M. Tomczak, Michał Talma, Stanley Brul

## Abstract

Since December 2019 a novel a coronavirus identified as SARS-CoV-2 or COV2 has been spreading around the world. On the 16th of May around 4.5 million people got infected and over 300,000 died due to the infection of COV2. The effective treatment remains a challenge. Targeted therapeutics are still under investigation. The papain-like protease (PL^Pro^) from the human SARS-CoV-2 coronavirus is a cysteine protease that plays a critical role in virus replication. Its activity is required to process the viral polyprotein into functional, mature subunits. Moreover, COV2 uses this enzyme to modulate the host’s immune system to its own benefit. Therefore, it represents a highly promising target for the development of antiviral drugs.

In this work, we discovered that ebselen, a synthetic organoselenium drug molecule with anti-inflammatory, anti-oxidant and cytoprotective activity in mammalian cells and cytotoxicity in lower organisms, is a highly active inhibitor of PL^Pro^CoV2. We proved that ebselen is a covalent, fast-binding inhibitor of PL^Pro^CoV2 exhibiting a low micromolar potency. Furthermore, we identified a difference between PL^Pro^ from SARS-CoV-1 (the corona virus which caused the 2002–2004 outbreak, SARS) and SARS-CoV-2 that allows to explain the difference in dynamics of the replication, and, thus, the disease progression. Namely, we present that they show differences in the binding affinity of substrates that we observed through kinetics and molecular docking studies. Using a novel Approximate Bayesian Computation method we were able to find kinetic constants for both enzymes. Molecular modeling study on the structure of the active site and binding mode of the ebselen with SARS and COV2 showed also significant differences that could explain our observation that ebselen is less active and slower bounding with SARS than COV2.

In conclusion, we show that ebselen inhibits the activity of the essential viral enzyme papain-like protease (PLpro) from SARS-COV-2 in low micromolar range.

## Introduction

More than 16 years after the global pandemic due to the SARS (severe acute respiratory syndrome) coronavirus (CoV), no anti-coronaviral medications have been developed for the treatment of infection caused by human coronaviruses (HCoV). SARS-CoV-1 (SARS) was identified as the causative agent of the fatal global outbreak of respiratory disease in humans during 2002–2003 and resulted in 8,422 cases with a case fatality rate (CFR) of 11% [1]. In September 2012 another SARS-like respiratory virus (termed Middle East respiratory syndrome coronavirus, MERS-CoV) had been established. MERS coronavirus spread around the world with a total of 178 laboratory-confirmed cases, but a CFR of as much as 43% [2] showing us how fatal the interspecies transmission potential of CoVs can be.

In December 2019, a novel coronavirus severe acute respiratory syndrome coronavirus 2 (SARS-CoV-2, CoV2) formerly known as the 2019 novel coronavirus (2019-nCoV), was discovered in Wuhan, China and was sequenced and isolated by January 2020 [4]. The disease, now termed coronavirus disease 19 (COVID-19), rapidly spread within China and infected a much larger number of people causing world economic and social paralysis. On May 16, 2020 nearly 4.5 million laboratory-confirmed infections were reported around the world, including over 300,000 deaths [5]. SARS-CoV-1 and SARS-CoV-2 are closely related, with studies highlighting that SARS-CoV-2 genes share >80% nucleotide identity and 89.10% nucleotide similarity with SARS-CoV genes [4].

The development of anti-coronaviral drugs remains challenging although a number of coronaviral proteins have been identified as potential drug targets [6,7]. Two of the most promising are Papain-like protease (PL^Pro^) and Main protease (M^pro^, also known as chymotrypsin-like protease 3CL^pro^) [6–8]. These proteases play an essential role in polypeptide processing during virus replication. PL^pro^, in addition to being crucial during replication via processing of the viral polyprotein [9], is proposed to be a key enzyme in the sustained pathogenesis of SARS-CoV. This includes deubiquitination [10] (the removal of ubiquitin), and deISGylation [11] (the removal of ISG15) from host-cell proteins. These last two enzymatic activities result in the antagonism of the host antiviral innate immune response [12]. As a result, PL^pro^ is an important potential target for antiviral drugs that may inhibit viral replication and weaken dysregulation of signalling cascades in infected cells that may lead to cell death in surrounding, uninfected cells [8].

Very recent studies led by Dikic confirmed the PL^Pro^ from SARS-CoV-2 (PL^Pro^CoV2) to be an essential viral enzyme and potential weak spot [13]. They proposed PL^pro^ to be Achilles’ heel of SARS-CoV-2. PL^Pro^ from SARS-CoV-1 (PL^Pro^SARS) and PL^Pro^CoV2 are closely related, with 82.9% sequence identity, and relatively distant from PL^Pro^ from MERS (32.9% identity). The removal of ubiquitin has been also recently confirmed by Rut et al. in [14].

Ebselen is a low-molecular-weight organoselenium drug that shows pleiotropic mode of action and due to its very low toxicity there are no barriers to using it in humans [15]. It is a well-known agent with therapeutic activity in neurological disorders [16] and cancers [17]. It also showed an antiviral effect on neurotropic viruses [18] and hepatitis C virus [19]. In a very recent work ebselen has been shown to attenuate inflammation and promote microbiome recovery in mice after antibiotic treatment for CDIAs [20]. Another recently published work proposed via virtual screening ebselen as a possible inhibitor of M^pro^ from SARS-CoV-2 [21].

Several lines of evidence demonstrated the biological effects of ebselen is mainly due to its antioxidant properties and capability of forming selenenyl-sulfide bonds with the cysteine residues in proteins [16,22–25].

Here, we demonstrate that ebselen inhibits activity of the essential viral enzyme, namely, papain-like protease (PL^pro^) from SARS-CoV-2 (PL^pro^CoV2) in low micromolar range. Moreover, we have identified the mechanism of inhibition as fast and irreversible as well as propose the binding mode of ebselen by molecular docking. We have found a difference in the mechanism of catalysis, the inhibition and the active sites between PL^Pro^ from SARS-CoV-1 and SARS-CoV-2. Furthermore, we used the recently published Approximate Bayesian Computation (ABC) methodology [26] to find kinetic parameters of the catalysis of Ubiquitin conjugated with fluorophore by PL^pro^SARS and PL^pro^CoV2. Our findings help to understand differences between SARS-CoV-1 and SARS-CoV-2 by analysing PL^pro^, and further indicate that ebselen is a highly active inhibitor of PL^Pro^CoV2 and indeed is a potential drug against COVID-19.

## Results

Following the high potential of ebselen as a promising drug, we sought to test its efficacy in the inhibition of the enzyme that is crucial in viral replication, namely, PL^pro^. Deep analysis of the active site of and mechanism of catalysis of coronaviruses PL^pro^ led us to conclude that small molecules with planar phenyl moieties, which also are able to modify cysteine residue active site, could be effective (Figure 1).

**Figure 1.**
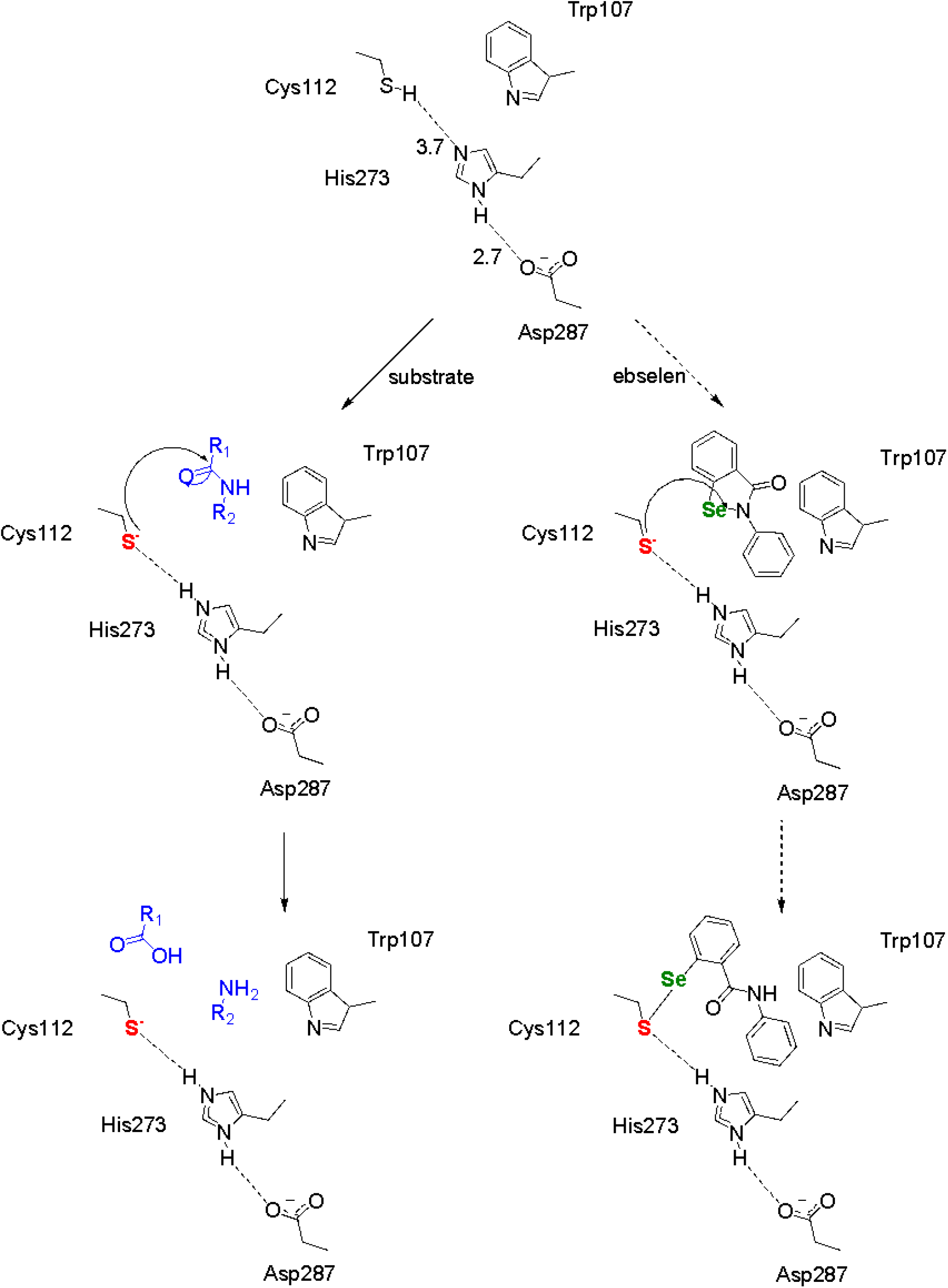
Analysis of the active site and mechanism of the catalysis (left) and possible inhibition by ebselen (right) of coronaviruses PL^Pro^ based on the active site of PL^Pro^SARS [26].

PL^pro^ from coronaviruses belong to the peptidase clan CA (family C16). The active site contains a classic catalytic triad composed of Cys–His–Asp. We analysed the active site and the mechanism based on the crystal structure of PL^pro^ from SARS-CoV-1 published by Báez-Santos et al. in [8,26] that showed PL^Pro^SARS has a catalytic triad composed of Cys112–His273–Asp287. This catalytic triad transforms the-SH group from cystine into a strong nucleophilic ion that attacks the carboxylic group of peptide bonds and leads to hydrolysis (Figure 1 (left)). The side chain sulfur atom of Cys112 is positioned 3.7 Å from the nitrogen in position 1 of the imidazole ring in catalytic histidine (His273). One of the oxygen atoms of the side chain of catalytic aspartic acid (Asp287) is located 2.7 Å from the nitrogen in position 3 of the same histidine (Figure 1). The side chain of Trp107 that is located within the oxyanion hole. The indole-ring nitrogen was proposed to participate in the stabilization of the negatively charged tetrahedral transition state of the reaction intermediates produced throughout catalysis [8,26,27].

Ebselen (Figure 1) meets active site requirements, size and conformation, and the ability to modify the-SH group. Moreover, ebselen possesses a clean safety profile in human clinical trials that in case of positive results can lead directly to discovering effective drugs against COVID-19. Encouraged by our analysis, we decided to test ebselen against both PL^Pro^ enzymes from SARS-CoV-1 and SARS-CoV-2.

As a substrate for our study we chose Ubiquitin conjugated with fluorophore (Ub-AMC). Progress curves (Figures 2A and 2C) at different levels of substrate concentration showed an interesting difference between two enzymes. PL^Pro^SARS catalyzed the reaction faster and achieved saturation. We estimated kinetic parameters of hydrolysis of Ub-AMC (Table 1) using the recently published novel Approximate Bayesian Computation (ABC) computational tool for calculating kinetic constants in the Michaelis–Menten equation [28]. This extremely useful framework gives us the opportunity to find the turnover number (*k*_cat_), the Michaelis Menten constant (*K*_M_) and, as a consequence, the catalytic efficiency of the enzyme (*k*_cat_/*K*_M_) without using high concentrations of Ub-AMC (Figures 2B and 2D).

**Figure 2.**
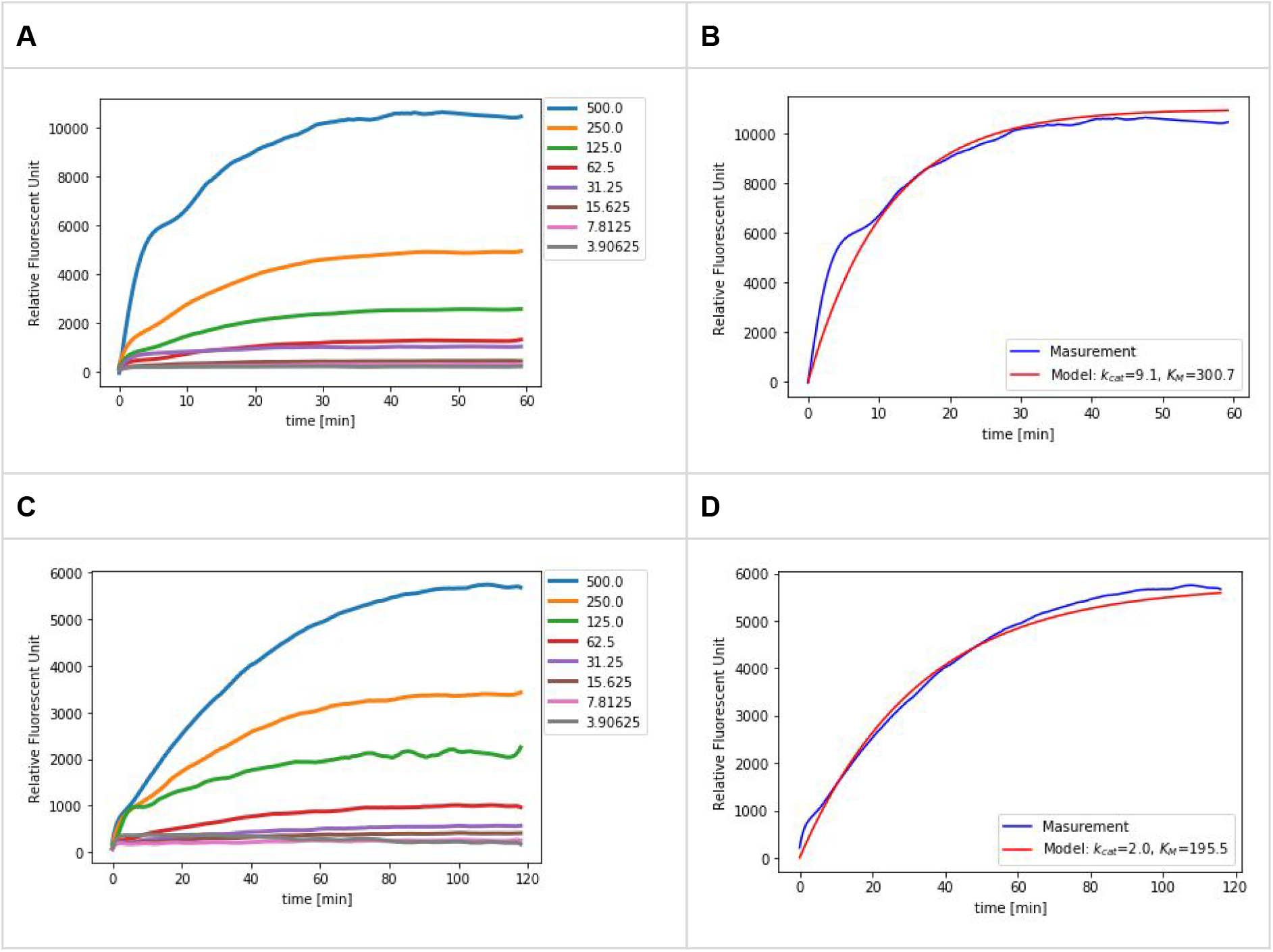
Progress curves for the hydrolysis of Ub-AMC by PL^pro^SARS (**A**) and PL^pro^CoV2 (**C**) (the concentrations of Ub-AMC are showed in the legend in nM, PL^Pro^SARS and PL^Pro^CoV2 were10 nM) and a comparison of measurements (in blue) and solutions of the Michaelis-Menten model for given parameters of enzymatic constants found by the ABC method (in red) for SARS-CoV-1 PL^pro^ (**B**) and SARS-CoV-2 PL^pro^ (**D**) at 500nM substrate concentration (*k*_cat_ and *K_M_* are expressed in s^-1^ and μM, respectively).

**Table 1.**
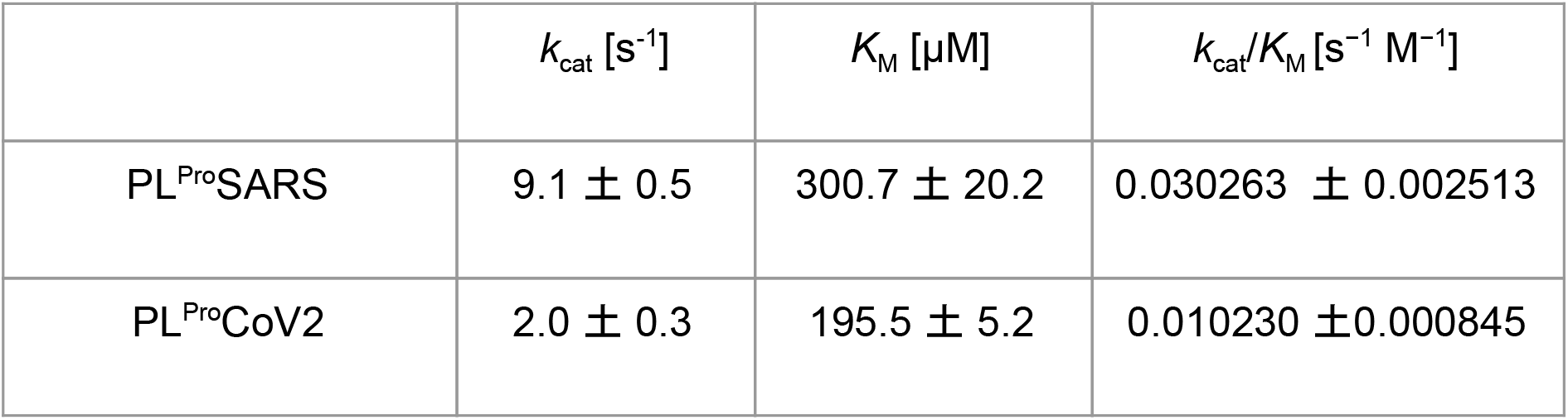
The kinetic data (*k*_cat_, *K*_M_, *k*_cat_/*K*_M_) for the Ub-AMC substrate of PL^Pro^SARS and PL^Pro^CoV2.

| | $k_{cat}$ [s<sup>-1</sup>] | $K_M$ [μM] | $k_{cat}/K_M$ [s<sup>-1</sup> M<sup>-1</sup>] |
| --- | --- | --- | --- |
| PL<sup>Pro</sup>SARS | 9.1 ± 0.5 | 300.7 ± 20.2 | 0.030263 ± 0.002513 |
| PL<sup>Pro</sup>CoV2 | 2.0 ± 0.3 | 195.5 ± 5.2 | 0.010230 ± 0.000845 |

The catalytic efficiency (*k*_cat_/*K*_M_) is often used as a specificity constant to compare the relative rates of reactions. Here we show that this ratio is three times higher for PL^Pro^SARS compared to PL^Pro^CoV2 that indicates its higher capability to hydrolyze Ub-AMC. PL^Pro^ is required for the processing of viral polypeptides and to modulate the host’s immune response, the higher efficiency may well contribute to the fact that once infected, SARS-CoV-1 was overall more aggressive and the disease developed faster.

We applied ebselen as a possible inhibitor and, indeed, it suppresses PL^Pro^ activity from CoV2 with inhibition constants approximately equal 2 μM (Table 1 and Figure 3). We determined the mechanism of inhibition of ebselen as irreversible, with steady-state binding being achieved immediately. Ebslen appeared to be an irreversible inhibitor of the studied PL^Pro^SARS as well, although in this case inhibition was weaker and the kinetics of binding was slow (Table 2). Irreversibility seems to confirm our first assumption that the inhibition of both enzymes can be associated with covalent bonds between Se from ebselen and S from cysteine. We confirmed irreversibility via dialysis and attemption of reactivation of the enzymes.

**Figure 3.**
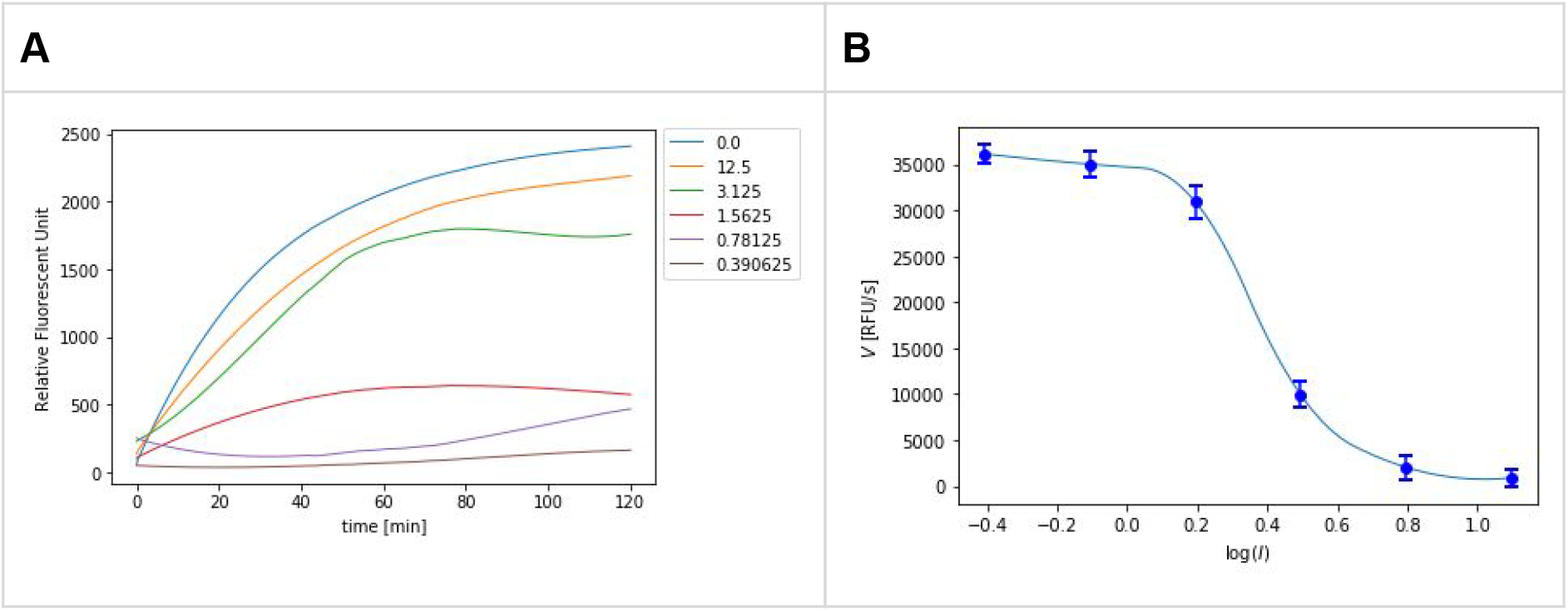
Progress curves for the hydrolysis of Ub-AMC by PL^Pro^CoV2 in the presence of increasing concentrations of ebselen (**A**) (the concentrations of Ub-AMC and PL^Pro^CoV2 were 50 nM and 10 nM, respectively, the concentration of ebselen are shown in the legend in μM). (**B**) A dose response curve showing the inhibition of PL^Pro^CoV2 by ebselen (the concentration of ebselen was from 12.5μM to 390nM).

**Table 2.** Inhibitory activities of ebselen toward two papain-like proteases from human coronaviruses SARS and CoV2.

| $IC_{50}[\mu M]$ | | | |
| --- | --- | --- | --- |
| PL<sup>Pro</sup>SARS |  | PL<sup>Pro</sup>CoV2 |  |
| no incubation | 60 min incubation | no incubation | 60 min incubation |
| >200 | $8.45 \pm 0.96$ | $2.02 \pm 1.02$ | $2.26 \pm 1.05$ |

The inhibitors were screened in TRIS buffer. The release of the fluorophore was monitored continuously. The linear portion of the progress curve was used to calculate the velocity. Each experiment was repeated at least three times and the results are presented as the average with standard deviation. For more details, please see the materials and methods section.

Our results were further illustrated by the use of molecular modeling to study the binding mode of ebselen with PL^Pro^SARS (Figure 4) and PL^Pro^CoV2 (Figure 4 and 5). This study confirmed our primary assumption that ebselen binds to the PL^Pro^CoV2 active site covalently and, thus, convinced us of our hypothesis about an irreversible mechanism of inhibition.

**Figure 4.**
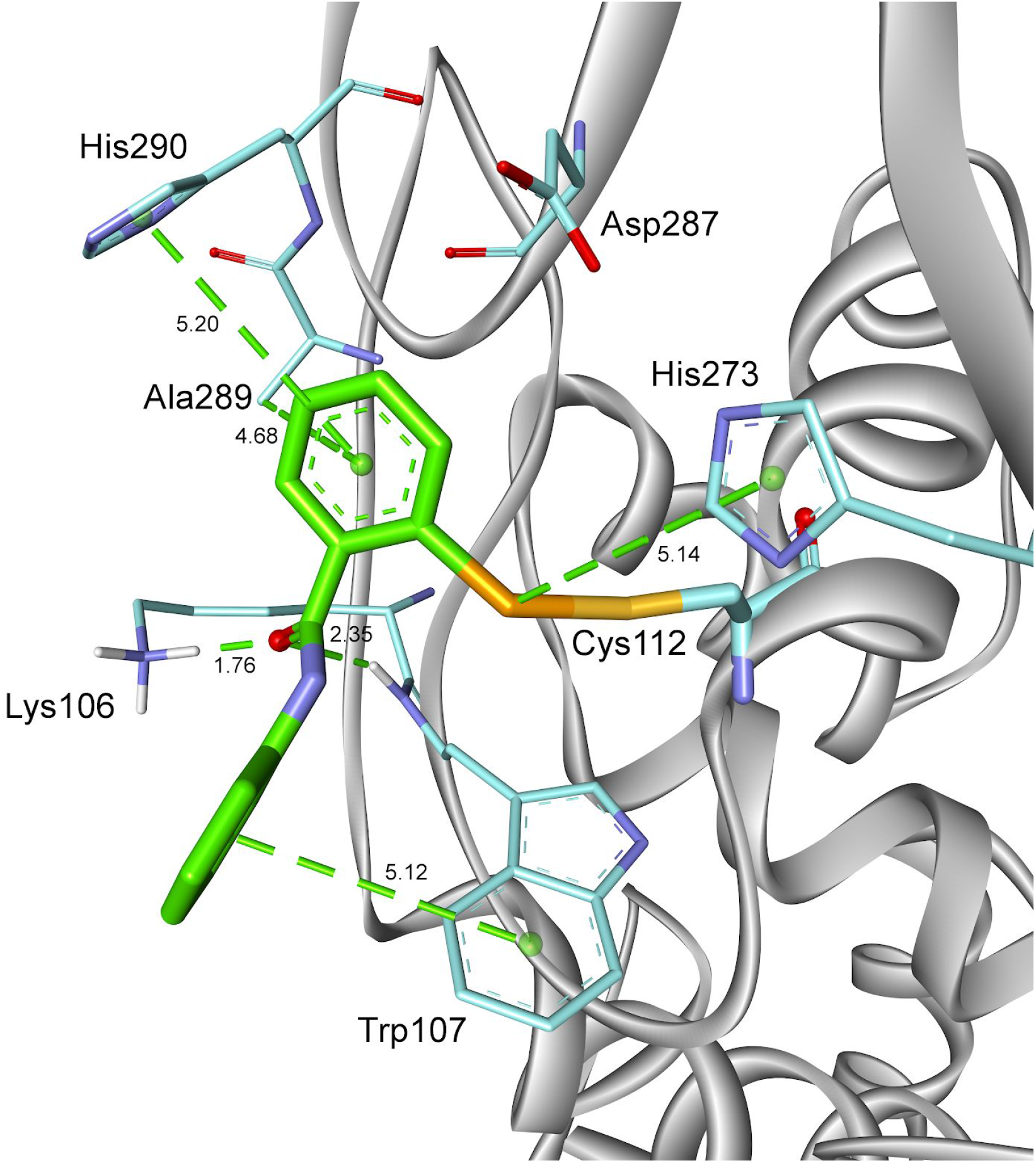
A model of the complex of ebselen with human SARS-CoV-1 PL^pro^ (PDB: 2FE8 [26]).

**Figure 5.**
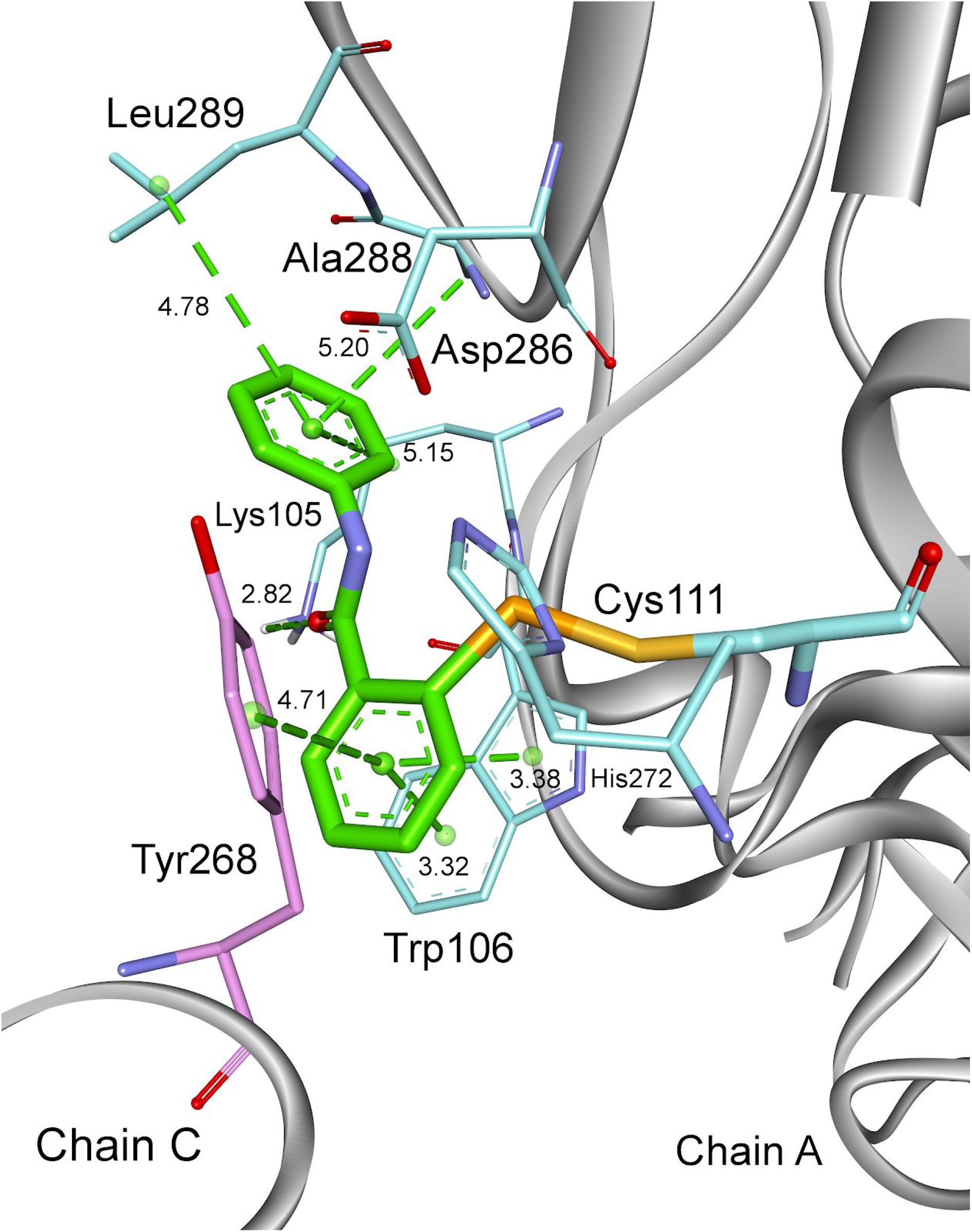
A model of the complex of ebselen with human SARS-CoV-2 PL^pro^ (PDB: 6W9C [29]).

In PL^Pro^SARS catalytic triad Cys112-His273-Asp287 is exposed externally out of the protein. The model with the most favorable thermodynamic stability shows that ebselen occupies an intersection between the putative catalytic triad Cys112-His273-Asp287 and Trp107, as we hypothesized. The *N*-phenyl ring of ebselen is surrounded by the aromatic ring of Trp107 (edge-to-face interaction) and Lys106. Amine group from Lys106 additionally interacts with oxygen from ebselen. Another edge-to-face interaction, we identified between *Se* -ring and an imidazole side chain of His290.

Ebselen is buried deeper in the active site of PL^pro^CoV2 than of PL^pro^SARS. This results from different conformation of the enzyme. In the PL^pro^CoV2 the catalytic triad Cys111-His272-Asp286 in each of three subunits are directed to the center of the protein. The most preferable conformation shows that the molecule occupies the same intersection between catalytic Cys111-His272-Asp286 triad and Trp106 but is additionally wrapped by other amino acids (Tyr268, Ala289, Leu298). Here, we identified a different binding mode where the *Se*-phenyl is directed to the oxyanion hole and Trp106, not *N*-phenyl as it was observed for PL^Pro^SARS. The *Se*-phenyl (from ebselen fragment) and an indole (from Cys322) formed π-π stacking interactions, while face-to-edge stacking interactions were observed with an aromatic ring from Tyr 268. The *N*-phenyl ring of ebselen adopts a bent-shaped conformation that fits well to the space between Leu298 and Ala288 forming π-alkyl interaction with them. The smaller distance between the oxyanion hole and ligand as well as more interactions with amino acids surrounding the active site in PL^pro^CoV2 could explain a better binding affinity observed in our experiments.

## Discussion

The present study provides an understanding of differences between SARS-CoV-1 and SARS-CoV-2 by analysing PL^pro^ and further highlights high potential of ebselen as a treatment for novel coronavirus appeared in December 2019. PL^pro^CoV2 is an essential viral enzyme that is required for the processing of viral polypeptides and assembling of new viral particles within human cells [13].

We first estimated parameters of kinetic constants and the catalytic efficiency of the catalysis of Ub-AMC by PL^pro^SARS and PL^pro^CoV2 (see Figure 2 and Table 1). Our results suggest that the capability to hydrolyze Ub-AMC is three times higher for PL^Pro^SARS compared to PL^Pro^CoV2. This observation is well-aligned with the fact that SARS-CoV-1 is more aggressive than SARS-CoV-2 and leads to a faster development of a disease.

Further, we showed that ebselen inhibits the enzyme PL^Pro^CoV2 and suppresses its activity with inhibition constants approximately equal 2 μM (see Figure 3 and Table 2). Moreover, we indicated that ebslen appeared to be an irreversible inhibitor of both PL^Pro^CoV2 and PL^Pro^SARS. However, it is weaker in the case of PL^Pro^SARS. Eventually, we studied the binding mode of ebselen with PL^Pro^SARS (Figure 4) and PL^Pro^CoV2 (Figure 5) with the use of molecular modeling. The obtained results firmly corroborated our primary assumption that ebselen binds to the PL^Pro^CoV2 active site covalently. This observation reinforces our view regarding the irreversibility of the mechanism of PL^Pro^CoV2 enzyme inhibition by ebselen and will aid in finding variants of the compound with further improved efficacy.

## Material and methods

### 1. General

Recombinant SARS-CoV-1 PL^pro^, SARS-CoV-2 PL^pro^ and Ubiquitin-AMC were purchased as 32, 11 and 250 μM solutions, respectively, from R&D Systems.

Ebselen (2-Phenyl-1,2-benzisoselenazol-3(2H)-one) is commercially available from Sigma Aldrich.

### 2. Enzymes assay

The enzymes were dissolved in a 50 mM Tris-HCl buffer containing DTT (2 mM), NaCl (5 mM) and 0.075% albumin, at pH 7.5, and preincubated 30 min. Spectrofluorimetric measurements were performed in a 96-well plate format working at two wavelengths: excitation at 355 nm and emission at 460 nm. The release of the fluorophore was monitored continuously at the enzyme concentration of 10 nM. The linear portion of the progress curve was used to calculate velocity of hydrolysis.

### 3. Inhibition assay

The inhibitor was screened against recombinant PL^pro^SARS and PL^pro^CoV2 at 37°C in the assay buffer as described above. For steady state measurement the enzymes were incubated for 60min at 37°C with an inhibitor before adding the substrate to the wells. Eight different inhibitor concentrations were used. Value of the concentration of the inhibitor that achieved 50% inhibition (*IC_50_*) was taken from the dependence of the hydrolysis velocity on the logarithm of the inhibitor concentration [I].

### 4. Molecular Modeling

Molecular modeling studies were performed using the Discovery Studio 2020 (Dassault Systemes BIOVIA Corp). The crystal structure of the SARS-CoV-1 and SARS-CoV-2 (PDB ID 2FE8 [26] and 6W9C [29], respectively) with protons added (assuming the protonation state of pH 7.5) was used as the starting point for calculations of the enzyme complexed with ebselen. The partial charges of all atoms were computed using the Momany-Rone algorithm. Minimization was performed using the Smart Minimizer algorithm and the CHARMm force field up to an energy change of 0.0 or RMS gradient of 0.01. Generalized Born model was applied. The nonbond radius was set to 14 Å.

## Acknowledgments

We gratefully acknowledge the Dassault Systemes for the free license for BIOVIA: Discovery Studio package given for our research.

EWT is co-financed by a grant Mobilność Plus V from the Polish Ministry of Science and Higher Education (Grant no. 1639/MOB/V/2017/0).

## Author contributions

EWT conceived the project. EWT designed the research and experiments with contributions from JT and SB. Experimental work was done by EWT. Parameter estimation was carried out by JT. Molecular docking was done by JT, EWT and MT. EWT, JT and SB drafted and revised the manuscript..

